# A versatile mitochondria isolation- and analysis-pipeline generates 3D nano-topographies and mechano-physical surface maps of single organelles

**DOI:** 10.1101/2021.10.31.466655

**Authors:** Saurabh Joshi, Friederike Hater, Jürgen Eirich, Joakim Palovaara, Henrik Ellinghaus, Paulina Heinkow, Hannah Callenius, Annette Peter, Ole Schweser, Martin Kubitschke, Murali Krishna Madduri, Amal John Mathew, Lucio Colombi Ciacchi, Janine Kirstein, Kathrin Maedler, Olivia Andrea Masseck, Iris Finkemeier, Manfred Radmacher, Rita Groß-Hardt

**Affiliations:** University of Bremen, Centre for Biomolecular Interactions, 28359 Bremen, Germany; Plant Physiology, Institute of Plant Biology and Biotechnology, University of Muenster, Schlossplatz 7, DE-48149 Muenster, Germany; University of Bremen, Institute of Biophysics, Otto-Hahn Allee 1, D-28359 Bremen, Germany; University of Bremen, Hybrid Materials Interfaces Group, Faculty of Production Engineering and Centre for Environmental Research and Sustainable Technology (UFT), 28359 Bremen, Germany

## Abstract

Living eukaryotic cells typically contain large quantities of highly dynamic mitochondria, which sustain the cells’ energy and redox homeostasis. Growing evidence suggests that mitochondria can functionally differ among but also within cells. The extent and biological significance of mitochondrial diversity is still largely unexplored, due to technical limitations that hamper profiling of individual organelles. Previous measurements of the cell’s interior have shown that membrane-bound compartments respond to metabolic manipulation by changes in their surface stiffness, suggesting that mechano-physical properties are a valuable readout of mitochondrial function. We here present the establishment of a robust multi-step analysis pipeline that allows one to profile mechano-physical properties of single mitochondria at the nanoscale using Atomic Force Microscopy (AFM). Firstly, we developed a **ra**pid **ce**ll-type specific isolation protocol (mRACE), which selectively functionalizes mitochondria with biotin, facilitating isolation by streptavidin decorated microbeads. We established the technique for human and rat cell cultures, the invertebrate *Caenorhabditis elegans*, and the model plant *Arabidopsis thaliana*. Based on this versatile tool, we detected diversity of mitochondrially associated proteins among different tissues, reflecting the trophic condition of the source material. Secondly, a rapid filtration-based mitochondria isolation protocol was established, which was combined with mRACE. Lastly, we established an AFM analysis platform, which generates 3D maps of the nano-topography and mechano-physical properties of individual mitochondria. The comparison of mitochondria with each other revealed an unprecedented diversity in their mechano-physical properties and suggests that shape is not the sole determining parameter for outer membrane stiffness. We expect our results to not only introduce a new dimension for basic mitochondrial research, but in addition to open the door for the exploitation of individual mitochondria for diagnostic characterization.

## Introduction

The cells of multicellular organisms are powered by mitochondria, which in addition to their function in ATP generation regulate important cellular processes including signal transduction, redox homeostasis and programmed cell death (1). A single cell typically contains between 100 and several thousand mitochondria (2), which, depending on the organism and cell type, can exhibit fascinating dynamics: Mitochondria can be highly mobile with respect to their position in the cell and travel along molecular highways (3, 4). In addition, the number and size of mitochondria are constantly changing due to mitochondrial fission and fusion (5-7). Previous research has mainly focused on the function of mitochondria as part of rather homogeneous populations. However, there has been a growing understanding that mitochondria differ between and even within cells and that these differences are biologically and pathophysiologically relevant (1, 8-10). Previous methodologies to study such diversity were limited as they largely relied on the physical separation of tissues and cells. Only more recent approaches have provided strategies to tag mitochondria of selected organisms in a cell-type specific manner with fluorophores (11) or fluorophore-tagged epitopes to enable affinity purification of the organelles from target tissues (12-14).

We here exploit a form-function correlation for the analysis of mitochondria, that capitalizes on the idiosyncrasy of the mitochondrial membrane structure. Mitochondria are structurally characterized by an outer and an inner mitochondrial membrane (MOM and IMM) with the IMM containing membrane invaginations termed cristae. The IMM is linked to ATP generation and contains the components of an electron-transfer chain that pumps protons against a gradient across the membrane. This electrochemical potential is used by the ATP synthase, a rotary nano-machine that generates ATP from ADP (15). It hence appears intuitive that the surface area of the inner mitochondria membrane correlates with the capacity of ATP production. In fact, a comparison of muscle mitochondria in recreationally active subjects and endurance-trained athletes indicates that cristae density is a better predictor of maximal oxygen uptake rate than mitochondrial volume (16). This structure-function correlation has also been described at the subcellular level. For example, a subset of mitochondria in the presynaptic membrane near active zones of the Calyx of Held, which is a large nerve terminal mediating early stages of sound localization, exhibit an increased ratio of cristae to outer membrane surface area indicative of high metabolic activity (17). Interestingly, it was previously shown that intracellular compartments including mitochondria can respond to metabolic manipulation by changes in their surface stiffness (18), suggesting that mechano-physical properties reflect metabolic activity.

What has thus far been lacking, however, is a technology which allows to literally get in touch with individual mitochondria for high-resolution mechano-physical profiling. To enable a characterization of individual mitochondria by Atomic Force Microscopy (AFM), we firstly developed a technique for cell-type specific isolation of mitochondria, analogous to a protocol for nuclei isolation (19) and similarly to a previously published protocol (12). We established the technique for vertebrates (human, rat), the invertebrate *C. elegans*, and the plant model *A. thaliana*. In addition, we established a platform for mitochondria chip-immobilization and high-resolution AFM profiling of individual organelles.

## Results and Discussion

### mRACE enables microbead- and chip-based isolation of mitochondria

There has been a growing understanding that mitochondria differ between and within cells and that these differences are biologically and pathophysiologically relevant. In order to study single mitochondria, we developed the mRACE protocol for the isolation of mitochondria from selected cell types. The method capitalizes on a multimodular mitochondrial targeted fusion protein (MTF), which combines three functions (Fig. 1A). First, the mitochondria-targeting domain of mtOM64, which has been shown to be sufficient to target fusion proteins to the MOM in *A. thaliana* (20-22). Second, a green fluorescence protein (GFP), which enables visualization of the tagged mitochondria in target cells and allows for live-cell imaging of mitochondrial dynamics. And third, a biotin ligase recognition site (BLRP), which provides a target site for biotinylation. The system is complemented by a second construct encoding the biotin ligase BirA, which attaches biotin to the BLRP acceptor site such that engineered mitochondria can be isolated by a biotin-streptavidin interaction. We aimed to establish mRACE for the isolation, enrichment and/or immobilization of mitochondria using streptavidin coated microbeads or streptavidin coated chips (Fig. 1F). Live cell imaging of MTF positive plants revealed dynamically moving and interacting fluorescent organelles (Movie S1). Closer inspection detected a fluorescence signal surrounding MitoTracker™ positive organelles, indicating that the chimeric protein was properly localized to the outer membrane of mitochondria (Fig. 1B). As there is substantial conservation of the anchor domains of MOM proteins in eukaryotic organism (23-25), we asked whether MTF could similarly be used to tag mitochondria in *C. elegans, human embryonic kidney* (HEK) *cells* and rat β-cells (INS-1E). Co-staining experiments with MitoTracker™ Red CMXRos or anti-TOM20-antibody revealed a colocalization with MTF in all other tested model organism (Fig. 1C-E), demonstrating broad versatility of the system for individuals of different kingdoms.

**Figure 1:**
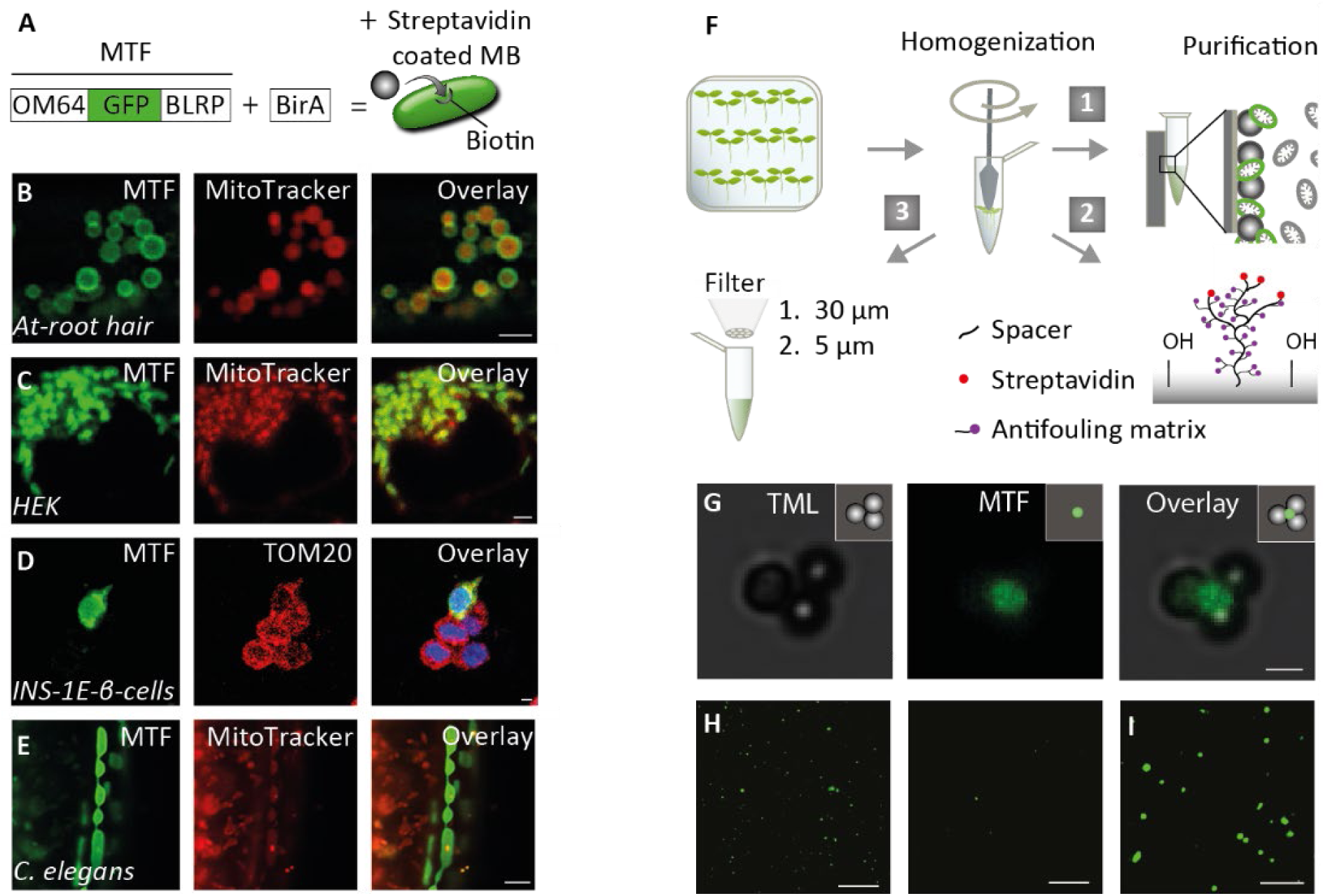
Exploiting mRACE for isolation and analysis of mitochondria from specific cell types. (A) The system capitalizes on the chimeric protein MTF, which contains the mitochondria targeting sequencing OM64, a GFP, and a biotin acceptor site. The latter gets biotinylated in the presence of the biotin ligase BirA. (B-E) mRACE-based mitochondria tagging in organisms of different kingdoms. MTF was overexpressed and co-expression of GFP and MitoTracker™ Red CMXRos or TOM20 is shown in single mitochondria of *A. thaliana* (B), of the human embryonic kidney cell line (C), in the rat insulin producing β-cell line INS-1 with DAPI-stained nuclei in the overlay (D; an example of five cells is shown, of which only one cell overexpressed MTF, while TOM20 remained stable in all cells), and *C. elegans* (E). (F) F1. Isolation and purification protocol recovering GFP-tagged microbead-bound plant mitochondria through a magnet. F2. Mitochondria-on-chip approaches: a sample recovered following homogenization and centrifugation can also be immobilized using a streptavidin functionalized chip surface in order to isolate and enrich for biotin labelled mitochondria. F3. For nonspecific immobilization, a sample recovered following homogenization can be subjected to two sequential filtering steps 30 and 5 µm followed by centrifugation on a chip to obtain individual mitochondria. (G) Transmission light, fluorescence light and overlay channels showing bead bound mitochondria isolated from *Arabidopsis* roots expressing *MTF* and *BirA* using strategy F1. (H) Specific enrichment of mitochondria immobilized on chips using strategy F2 showing MTF and BirA expressing samples (left) over samples only expressing MTF (right). (I) Mitochondria isolated using strategy F3 and immobilized on chips using centrifugation. Scale bar 2 µm (B-E), 1 µm (G), 10 µm (H, I).

To investigate whether mRACE can be used to enrich and isolate mitochondria for various downstream applications, we generated *Arabidopsis* plants expressing *MTF* and *BirA* under the broadly active *p35S* promoter (Fig. 1F). We isolated mitochondria using streptavidin-coated microbeads based on a previously established protocol (12) (Fig. 1F1) and recovered bead-bound GFP-tagged mitochondria (Fig. 1G).

In order to bind mitochondria to streptavidin-coated chip surfaces, we adapted the protocol to rapidly isolate and enrich mitochondria (Fig. 1F2). Microscopic inspection documented specific binding of MTF functionalized mitochondria from BirA expressing over non-BirA expressing *Arabidopsis* plant root samples (Fig. 1H). Additionally, we established a novel protocol that allows rapid isolation of mitochondria in ten minutes. The approach is based on filtering the fluorescently labelled mitochondria sequentially using 30 µm and 5 µm diameter cell strainers from a crude cell lysate (Fig. 1F3). Depending on the prevalence of the cell type under investigation, mRACE also allows to chip-immobilize mitochondria by centrifugation and subsequent selection of target mitochondria based on their fluorescence (Fig. 1I).

In the following, we outline procedures demonstrating the broad versatility of this methodology for downstream biochemical and biophysical applications.

### mRACE efficiently enriches the mitochondrial signatures of source tissues

To investigate whether and to what extent mRACE-isolated mitochondria reveal mitochondrial diversity among different tissue types, we subjected *Arabidopsis* root and seedling-derived samples to an LC-MS/MS analysis. We identified 6265 protein groups throughout all runs, including those identified in the pre-bead lysates (Fig. S1, Table S1), of which 688 protein groups were mitochondrial according to the SUBA4 database (26). Compared to control samples that did not contain the mRACE constructs, 528 mitochondrial protein groups were significantly enriched in the pulldowns of at least one of the tissue types (log_2_FC > 1, adjusted p value < 0.05 (LIMMA), quantified in all 3 replicates).

For the seedling sample, intermediate isolation steps were also analyzed. Both, the pre-bead sample and the supernatant revealed an even distribution of mitochondrial and non-mitochondrial proteins (Fig. S1), indicating that the combination of biotin functionalized mitochondria and streptavidin decorated microbeads were causative for the selective enrichment of mitochondrial proteins. Importantly, GO assignment for the mitochondrial proteins in the mRACE samples detected proteins from all mitochondrial sub-compartments, indicating that the isolation procedures allowed to recover intact mitochondria (Fig. 2B). The SUBAcon localization of all identified proteins shows that apart from mitochondrial proteins, proteins associated with the Golgi apparatus are enriched in both seedling and root samples (Fig. 2A). Plastid-associated proteins are also slightly enriched in both samples with enrichment being more pronounced in the photosynthetic active seedling tissue than the root sample (Fig. 2A, right). Together, these data show that mRACE is a powerful tool for the robust enrichment of mitochondrial proteins from intact mitochondria (Fig. 2A-B).

**Figure 2:**
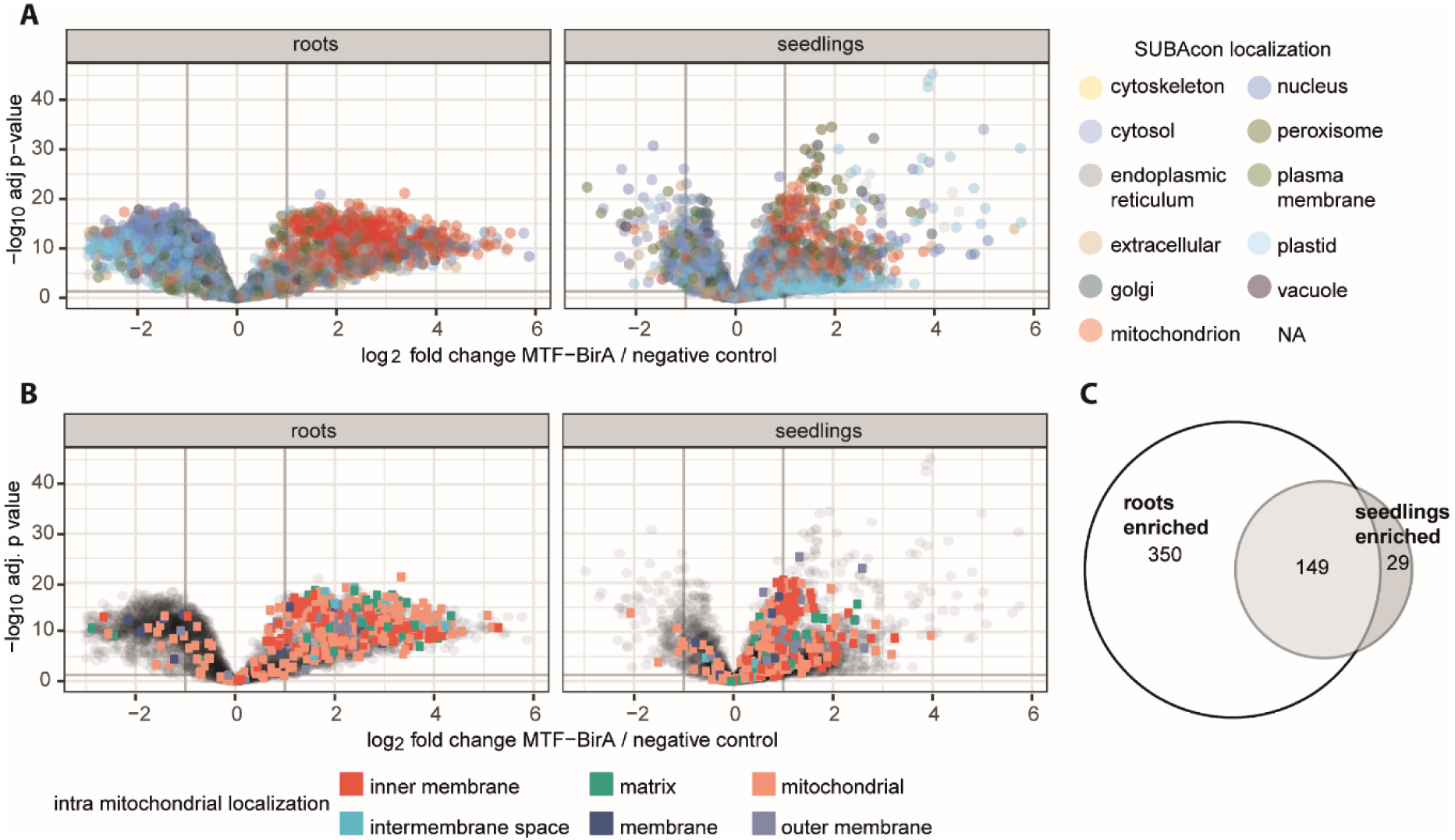
Proteome comparison of mRACE isolated mitochondria from photosynthetic and heterotrophic tissues. Mitochondria were isolated from seedlings (photosynthetic active tissue) and root (heterotrophic tissue) cultures using mRACE. Seedlings and root cultures not carrying the mRACE constructs were used as control. All samples were subjected to LC-MS/MS analysis. (A) Volcano plot comparing mRACE-isolated root and seedling samples to control. (B) Volcano plot comparing designated localization of mitochondrial proteins between mRACE-isolated root and seedling samples to control. (C) Venn diagram illustrating the different sets of mitochondrial proteins that are either exclusive or shared between roots and seedlings (149 overlapping IDs). All experiments were conducted in biological triplicates.

### mRACE detects tissue-specific mitochondrial diversity reflecting the trophic condition of the source tissue

When comparing mitochondrial proteins in seedlings (photosynthetic active tissue) and roots (heterotrophic tissue), we found 29 proteins specifically enriched in photosynthetic tissue and 350 proteins enriched in heterotrophic tissue, with 149 proteins being enriched in both tissue types (Fig. 2C, filters applied are log_2_FC > 1, adjusted p value (LIMMA) < 0.05, quantified in all 3 replicates). In order to identify tissue-specific differences in the isolated mitochondria, we calculated an enrichment score (log_2_FC x –log_10_(p value)) for every mitochondrial protein per tissue. Table S1 contains a list of all detected mitochondrial proteins according to SUBAcon and their respective values. In addition, we calculated the score difference as a measure for tissue-specific enrichment and subjected this ranked list to a Functional Enrichment Analysis via the String DB (27). The respective GO terms for biological processes and molecular functions are provided in Fig. S2. As expected, mitochondria isolated from roots (Fig. S2) were richer in proteins associated with aerobic respiration and oxidoreductase activity, while mitochondria from seedlings were enriched in RNA binding proteins. The most striking difference between seedling and root tissue is FORMATE DEHYDROGENASE (FDH, At5g14780), a protein involved in stress adaptation (28-31), which ranks among the six most abundant proteins in mitochondria from root tissue, while in seedling it ranks only among the 247^th^ most abundant proteins (Table S1). This observation is different from previous reports, where FDH was reported to be significantly enriched in photosynthetically active tissue (32, 33) and might be accounted for by the different root cultivation regimes.

### A novel AFM-based analysis platform generates a high-resolution 3D nano-architecture of individual mitochondria

Protein analysis indicates that mitochondria stay intact during the isolation procedure and we next asked whether chip immobilized mitochondria were still active. To detect metabolic activity of individual isolated mitochondria, we used a mitochondrially targeted pH sensor (34) that monitors the physiological status with single mitochondrion resolution. cpYFP expressing tissue homogenates were subjected to homogenisation and filtration (Fig. 1F3) and immobilized on coverslips using a previously established protocol (34). Differential treatment of chip-immobilized mitochondria resulted in differential fluorescence intensities (Fig. 3A). Mitochondria supplemented with succinate, ATP and rotenone gave a strong and dynamic fluorescent signal as compared to mock-treated mitochondria, indicating that the organelles were metabolically responsive. Interestingly, some mitochondria displayed short transient pulses while others displayed stronger and longer pulses. Future experiments will help to elucidate the prevalence and functional significance of this differential responsiveness to electron transport chain effectors.

**Figure 3:**
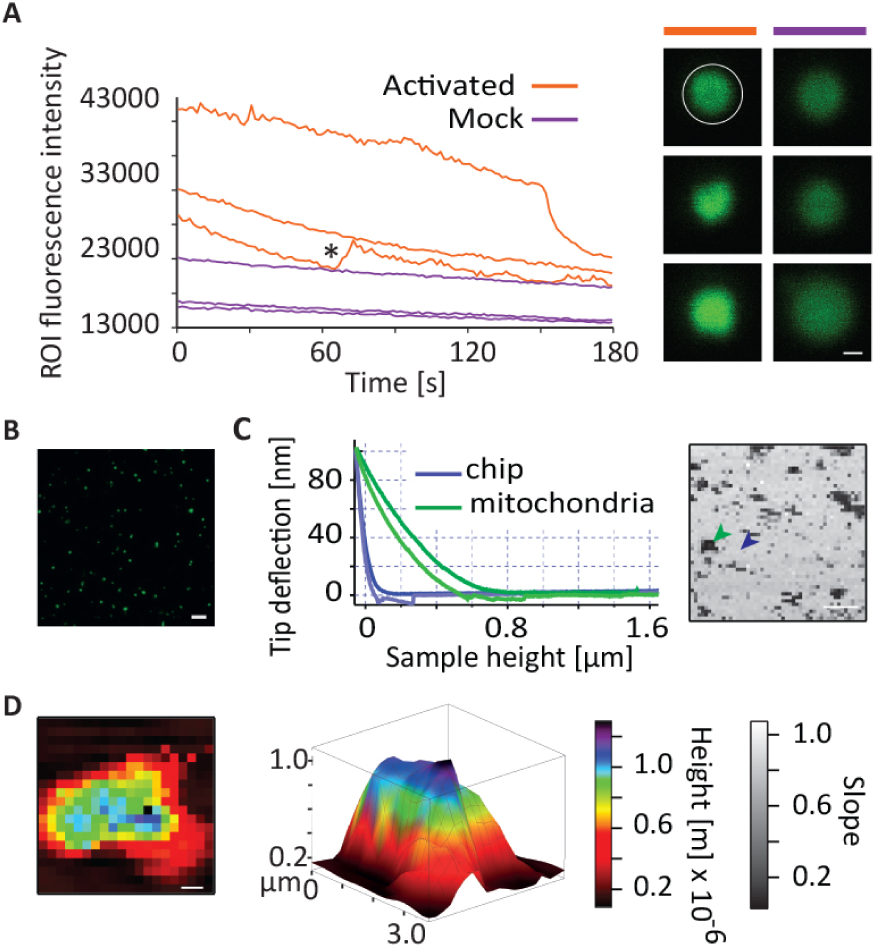
Characterising the 3D-topography and mechano-physical properties of single mitochondria: (A) The right panels show cpYFP signals of individual mitochondria upon activation (10 mM Succinate, 0.25 mM ATP and 20 µM rotenone) and mock ‘basal incubation medium’ treatment. The left panel shows quantification of the fluorescence intensities of chip immobilized cpYFP labelled individual mitochondria in the region of interest (white circle at right) of 3 activated and 3 mock treated individual mitochondria. * Single mitochondrion showing a temporal increase in fluorescence intensity at 72 secs. (B) Application of GFP labelled mitochondria on a chip facilitates physical organelle separation making them accessible to AFM studies. (C) Graph showing approach (darker shade) and retract force curves representing the arrowhead marked areas on the force map shown to the right. D) The height of the mitochondrion was calculated on the basis of the contact point determined in each force curve and a 3D reconstruction of the mitochondrion was generated. Scale bar 0.2 µm (A), 10 µm (B, C), 0.5 µm (D).

To introduce a new dimension of analysis, we aimed to make individual mitochondria amenable for the analysis of mechano-physical properties and subjected chip-immobilized mitochondria to Atomic Force Microscopy (Fig. 3B). A force map with a resolution of 50 × 50 force points on a 50 µm^2^ was obtained, in a first step, to locate mitochondria (Fig 3C). The slope of the approach curve recovered on a stiff substrate was close to 1, while values of around 0.1 were indicative of biological material (Fig 3C). Based on the contact point information, 3D reconstructions of the mitochondrion’s height were generated displaying the mitochondrial architecture. In contact mode imaging, height is determined at a positive loading force, which can generate slightly compressed representations in case of soft samples. In order to avoid this problem and to reflect the true topography, we here derived the height information from the contact point, where the loading force is zero (Fig. 3D).

### AFM-based profiling of individual mitochondria uncovers an unprecedented mechano-physical diversity between individual mitochondria

We further optimized the system by combining the AFM with fluorescence microscopy, which allowed to unambiguously identify single mitochondria on chips. Using DirectOverlay imaging, mechano-physical data obtained on three individual mitochondria were superimposed on their fluorescence image confirming data acquisition on mitochondria (Fig. 4A).

**Figure 4:**
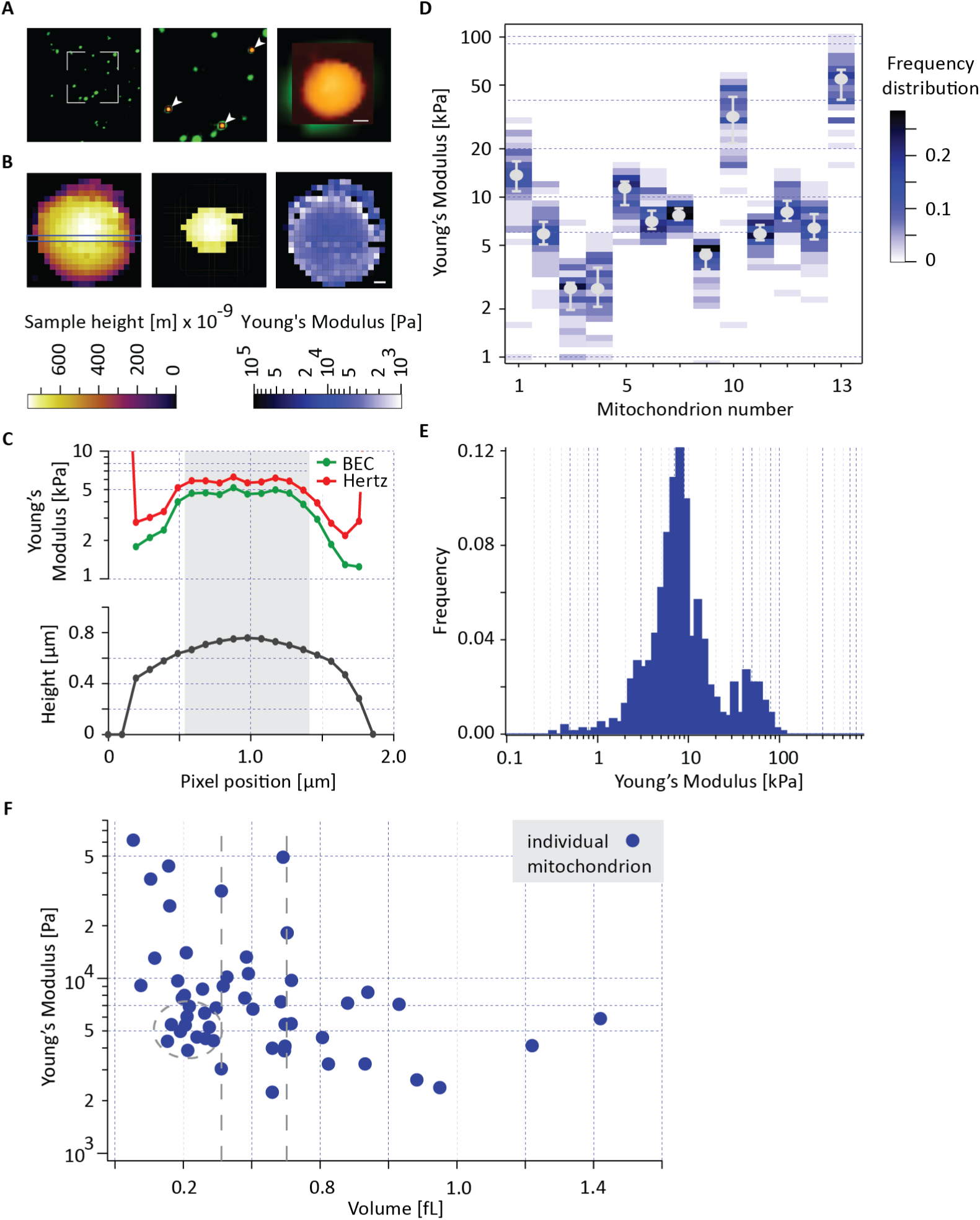
AFM reveals an unprecedented diversity in mechano-physical properties within and among individual mitochondria: (A, B) Young’s Modulus values on individual mitochondria. (A) On an area of choice (white edges), force maps were measured on individual mitochondria (arrow heads) obtaining a direct overlay image of fluorescence and biophysical information. (B) A typical sample height profile determined from the contact point of each individual force curve (left panel). Middle panel showing top 15% of the data points that are used for calculating the median as a measure for the representative YM of this mitochondrion. In right panel, YM profile generated on this mitochondrion using a combination of Hertz-Fit and BEC model. (C) YM and height along one line of a two-dimensional force map obtained from the blue rectangle in left panel B. Young’s moduli are determined by the conventional Hertz model (red) or the BEC model (green) which corrects for the bottom effect. Grey bar denotes data pixels included in top 15%. (D) Histogram profile showing YM variations within and among mitochondria. YM shown as median with whiskers representing 25^th^ and 75^th^ percentile. (E) Compiled histogram profile segregating mitochondria in two mechanical categories. (F) Graph plotting the YM as a function of the geometrical data measured by AFM. Grey dashed oval points to a cluster of mitochondria displaying similar mechanical and volume profile while dashed lines mark individual mitochondria with similar volume showing contrasting YM. Whiskers representing 25^th^ and 75^th^ percentile not shown in F for simplicity but shown in Fig. S4. Scale bar 0.2 μm (A,B)

The elastic or Young’s Modulus (YM) is a universal parameter reflecting stiffness of the sample. It is obtained by fitting the so called Hertzian model (35) to the force vs indentation data on each force curve (Fig. S3). To reduce artefacts we processed the data in two steps: First, we corrected the bottom effect, i.e. the fact that the AFM tip “feels” the underlying substrate, using a modification of the Hertz model derived by Garcia et al (36). Second, we integrated only those data points taken on the top of mitochondria (Fig. 4B, C) since data recovered at the periphery were unreliable due to the sample slope. On the basis of the processed data, we asked whether there are any similarities in the mechanical properties of individual mitochondria. We were able to measure 13 individual mitochondria recovered from an *Arabidopsis* plant root sample in high resolution. The median YM values on these mitochondria showed striking differences ranging from 2.7 kPa to 50 kPa (Fig. 4D), uncovering a previously unprecedented diversity among mitochondria with respect to their outer membrane stiffness. Interestingly, the histogram on extracted values on individual force maps, in addition, detected a striking variation in the YM values on individual mitochondria (Fig. 4D). A histogram over all YM values from 13 mitochondria suggested two different mitochondria categories showing peaks around 8 and 50 kPa, respectively (Fig. 4E). To gain a more comprehensive view, we plotted YM as a function of geometrical data for 52 individual mitochondria. We, combined height and circumference as a proxy volume for the typically spherically shaped isolated mitochondria (Fig. 4F). We observed a correlation where volume was indirectly proportional to YM and clusters of mitochondria that comply to this observation showing similar mechanical and volume profiles. Interestingly, we also detected mitochondria with similar volume profiles that showed striking YM differences. Our data thus suggest, that shape is not the sole determining factor for outer mitochondrial stiffness in support of the previous findings that have implicated mitochondrial activity as an important factor for the mechano-physical properties of the MOM (18).

Many disorders including diabetes, cancer, cardiovascular and neurodegenerative diseases have been associated with mitochondria dysfunction. Our findings open new avenues for exploiting mitochondria-on-chip approaches for the characterisation of mitochondrial 3D topographies and mechano-physical properties at high resolution. In light of previously described correlations between mechano-physical properties of mitochondria and their activity, we expect our results to also introduce a new dimension for functional analysis of single mitochondria. In addition, we propose that the analysis pipeline developed in this work can easily be adjusted to other organelles.

## Materials and methods

### Plant growth conditions

Seeds were sterilised by incubating them twice in 70 % EtOH for 2-4 minutes and two rinsing steps in 99,9 % EtOH. Sterilized and dried seeds were sown onto MS Medium (0.23 % (w/v) MS basal salt, 0.05 % (w/v) MES, 0.8 % Phyto Agar, pH 5.9 with KOH) containing 25 µg/ml Kanamycin and 25 µg/ml PPT if necessary, for selection. After stratification at 4°C for two days in darkness, germination took place at 23°C under long day conditions in an upright position.

Arabidopsis root cultures (ARC) were prepared as described by Czakó, Wilson (37) with modifications. Approx. 40 7-days-old seedlings were transferred to ARC-Medium (0.43 % (w/v) MS Basal Salt, 0.3 % (v/v) Miller’s solution (6 % (w/v) KH_2_PO_4_), 0.2 % (v/v) Vitamix Stock (0.5 % (w/v) Thiaminuteshydrochloride, 0.05 % (w/v) Pyridoxinhydrochloride, 0.1 % (w/v) glycine, 0.05 % (w/v) nicotic acid, 0.025 % (w/v) folic acid, 0.05 % (w/v) biotin), 0.02 % (w/v) myo-inositol, 3 % (w/v) sucrose) in a sterile 500 mL flask. Cultures were incubated at 25°C, 80 rpm, and in darkness. After three weeks medium was exchanged with fresh ARC-Medium. ARCs were used for mRACE after 5-6 weeks.

### Cloning and transformation

Both mRACE constructs were cloned on the basis of the INTACT constructs (19). The *NTF* was subcloned into a pGIIBar vector (38). For the MTF the first 138 bp of *mtOM64* (AT5g09420), encoding the mitochondrial targeting domain (20), were amplified from *Arabidopsis* gDNA with primers introducing a *Pac*I and a *Hin*dIII restriction site. The *eGFP* of the *NTF* was also amplified to introduce a *Hin*dIII restriction site at its 5’ end. The subcloned *pGIIBar_NTF* was restricted with *Pac*I and *Eco*RV, excising *WPP_eGFP*, and the newly amplified 138bp of *mtOM64* and *eGFP* were introduced into the construct. The *BirA* was subcloned into a pGIIKan vector (38). For both constructs *p35S* promoter was introduced via *Asc*I/*Pac*I restriction sites. *A. thaliana* var. Columbia plants were transformed using the floral dip method (39) and selected for transformants with either Kanamycin or Basta®.

Cloning details for molecular markers shown in Figure 1 panel C to E upon request.

### Mitochondria isolation with mRACE

#### mRACE mitochondrial isolation from ARC

All steps were performed at 4°C or on ice. Approx. 30 g of ARC were homogenized with 100 mL ice cold (ic) KPBS (136mM KCl, 10 mM KH_2_PO_4_, pH 7.25) in a mortar on ice. Homogenate was filtered through four layers of sterile Gauze swabs (Hartmann AG). Remaining plant material was again added to the mortar, re-homogenized with additional 50 mL of ic KPBS and filtered again through four layers of sterile Gauze swabs. Homogenate was centrifuged at 2000 x g for 5 minutes on 4°C. Supernatant was carefully transferred to a new tube and centrifuged at 16,000 x g for 10 minutes on 4°C. Mitochondria containing pellet was resuspended in 1 mL ic KPBS and added to 40 µL Dynabeads™ MyOne™ Streptavidin T1 (Thermo Fischer) in ic KPBS. Crude mitochondrial isolate was incubated with the beads at 10 rpm over-head-rotating, for 10 minutes on 4°C. Afterwards 1 mL ic KPBS was added and sample was placed on a magnet for 2 minutes. The supernatant was discarded, and the pellet was washed on the magnet with 1.2 mL, 800 µL, 500 µL, 300 µL and 100 µL ic KPBS successively. Finally, pellet was resuspended in 100 µL ic KPBS.

#### mRACE mitochondrial isolation from seedlings

All steps were performed at 4°C or on ice. 400 mg of 7 days old seedlings were homogenized with 300 µL ic KPBS using the homogeniser (Büchi). Homogenate was filtered through a 30 µm cell strainer (Sysmex). Homogenization tube was washed with 200 µL ic KPBS and remaining liquid was filtered into the homogenate as well. Homogenate was then centrifuged at 2500 x g for 5 min on 4°C. Supernatant was carefully transferred into a new tube and centrifuged at 15,000 x g for 15 minutes on 4°C. Pellet was resuspended in 100 µL KPBS and incubated together with 20 µL (Thermo Fischer) in KPBS at 10 rpm over-head-rotating, for 10 minutes at 4°C. Afterwards 800 µL ic KPBS were added and sample was placed on the magnet and incubated for 2 minutes at 4°C. Supernatant was discarded and pellet was washed on the magnet with 800 µL, 500 µL, 300 µL, 200 µL and 100 µL successively. Finally, the pellet was resuspended in 100 µL ic KPBS

#### mRACE mitochondrial specific immobilization

Mitochondrial isolation from approx. 40 mg of seedlings (∼14 days old) was performed as per mRACE for seedlings. Pellets were resuspended in 400 µL of KPBS buffer and 100 µL of sample was added on a pre-cut (with diamond knife) PolyAn covalently coated streptavidin chip. Samples were incubated at 4°C for 10 mins. Chips were washed 5x using 3 ml KPBS buffer in a 35 mm Petri plate on a shaker at 300 rpm for 1 min each. For imaging, chips were carefully placed on a µ-Dish ^35 mm, low^ (IBIDI GmbH) imaging dish containing KPBS buffer and images were captured using Leica DMI6000b epifluorescence inverted microscope equipped with GFP filter cube. Images (Fig. 1H) were equally processed in inbuilt Leica LAS AF software.

#### mRACE mitochondrial filtration and unspecific immobilization

Roots from ∼20 *A. thaliana* seedlings (∼14 days old) were harvested and collected in a 1.5 mL Eppendorf tube containing 400 µL of 1x KPBS buffer (40). The root tissues were crushed open using homogenizer (Büchi) to release organelles by mechanical lysis of cells for 2 mins. The crude cell lysate was filtered through a pore size of 30 µm diameter cell strainer (Sysmex). The collected flow through was filtered through a fine pore size of 5 µm diameter cell strainer (pluriSelect) by successive centrifugation steps with gradual increase in force (1 min at 60 x g, 30 s at 94 x g, 30 s at 211 x g, 1 min at 376 x g). All steps were performed at 4°C. Mitochondrial immobilization was adapted from (34). In brief, the subsequent filtrate containing GFP labelled mitochondria were immobilized on a clean round 24 mm diameter coverslip by centrifugation (800 x g for 5 mins at 4°C) in 1x KPBS buffer. Immobilized mitochondria (Fig. 1I, Fig. 4A) were imaged using epifluorescence microscope assembled with AFM. Fluorescence images were processed in inbuilt JPK data processing software.

### Mitochondria isolation using Density Gradient Centrifugation (DGC)

Isolation of mitochondria from ARC was adapted from a previously published protocol (41) using sucrose gradient solutions prepared with 20 %, 36 %, 40 %, and 5 2% (w/v) in a basic buffer containing 50 mM Tris, 6.3 mM EDTA and 0.1 % (w/v) BSA, pH 7.5. Samples were stored in 10 % glycerol solution at -80°C. DGC isolated mitochondria were used to measure mitochondrial topography.

### In-house chip activation and functionalization for mitochondrial immobilization

The cleaning and activation of the chip was performed according to method 3 (42). Silanization of the chip was done using 1kDa Biotin-PEG-Silane (Creative PEGWorks, Durham, USA) in vial containing absolute ethanol for 24 hrs. Upon washing with ethanol, streptavidin binding was facilitated by incubating the chip in a streptavidin solution containing PBS at 4°C for 1h. Functionalized chips were subjected to DGC isolated MTF harbouring biotinylated mitochondria to obtain mitochondrial immobilization.

### Atomic force microscopy

An MFP-3D AFM (Asylum Research, Santa Barbara, CA, USA) was used to study and sense the topography of immobilized mitochondria. The AFM head was placed on an inverted optical microscope (Zeiss Axiovert 135 TV) that allowed us to visualize the sample and AFM tip. A JPK Nanowizard 3 Ultra (Bruker) was used in this study to measure the mechanics of mitochondria. The AFM head was placed on an inverted epifluorescence microscope (Zeiss Axio Observer D.1) that allowed us to visualize samples and AFM tip. For fluorescence imaging the samples were illuminated using a lamp source X-Cite 120-Q series. The fluorescence image was captured using Andor Zyla sCOMS USB 3.0. A PeakForce QNM-Live Cell Calibrated (PFQNM-LC-CAL) probe (Bruker AFM Probes, France SAS) was used to image mitochondria. These pre-calibrated tips have a force constant of around 0.1 N/m with tip length 17 µm, tip radius 75 nm, opening angle 20°. For vibration and noise cancellation, the AFM set up was placed on a Halycionics_i4 Accurion granite slab. The entire set up was placed in a Bruker vibration and noise cancelling chamber.

### Data acquisition

For accurate force measurements, the deflection sensitivity was determined by the thermal tune method (43) before each experiment using the pre-calibrated force constant provided by the manufacturer. Data acquisition was performed in 1x KPBS buffer. The immobilized mitochondria on a 24 mm diameter coverslip were mounted on a coverslip holder placed on the AFM stage to position in x – y direction. The optical calibration was done using Direct Overlay feature to superimpose fluorescence and mechanics data. Topographic and mechanical data were obtained using contact mode. An array of force curves was collected over a desired area superimposing a GFP labelled mitochondrion with a resolution of 16 × 16 or 20 × 20 force points with speed 5 force curves/ sec at rate 20 kHz, typically over a field of view of 2 µm.

### Data analysis

The data acquired using the JPK Nanowizard were imported and analysed using a home built software written in IGOR (WaveMetrics, Lake Oswego, USA) (44). Force curves were fitted with the Hertz model (35) for a spherical indenter (radius of curvature 75nm) in a region of interest (between 150pm and 1.5nm deflection, corresponding to a force range of about 20-100 pN to yield Young’s modulus. We calculated sample height (thickness) from the contact point data of each force curve. First, all force curves taken on the substrate were determined by considering only points where the slope of the approach curve was larger than 0.1. A plane was fitted to these contact point values, and then this fit plane was subtracted from all contact point values to get the sample height. In a second fit, now the force curves were fitted with bottom effect correction derived by Garcia et al (36). To obtain a typical value of the Young’s modulus the median of all values on-top of the mitochondrion was calculated. “On-top” was defined as the data obtained on top 15% highest force curves (entire, not line by line) on mitochondrion starting from highest data point.

### Confocal microscopy

#### Colocalisation experiment Arabidopsis

Seven day old seedlings were incubated with 100 nM MitoTracker™ Red CMXRos (Thermo Fischer) in liquid MS Medium (0.23 % (w/v) MS basal salt, 0.05% (w/v) MES, pH 5.9 with KOH) for 30 minutes at 4°C in darkness. Afterwards seedlings were incubated in MS Medium only for 30 minutes at RT in darkness. Microscope slides were prepared by placing the seedlings in a drop of 10 % glycerol and covered with a cover slide. Pictures were taken using the Airyscan super resolution mode of the LSM 880 Indimo, AxioObserver (Zeiss) with the Plan-Apochromat 40x/1.4 Oil DIC M27 objective, the MBS 488/543 beam splitter and with excitation wavelength of 488 nm, 1.0%, and 543 nm, 1.0%. The pixel-dwell was 8.19 µs.

#### Colocalisation experiment in HEK cells

*Human embryonic kidney cells* were transiently transfected with MTF and costained with MitoTracker™ Red FM. Confocal pictures were taken at an Airyscan LSM880 (Zeiss) with an 40x Oil objective.

#### Colocalisation experiments in INS-1E-β-cells

The rat β-cell line INS-1E was transiently transfected with MTF for 48h using jetPRIME^®^ transfection reagent (#114-75; Polyplus transfection, France) according to manufacturer’s instructions; jetPRIME-plasmid-DNA complexes were added to complete RPMI-1640 medium. Efficient transfection was evaluated based on western blot and fluorescent microscopy.

Cells were fixed and costained with MitoTracker™ Red CMXRos (#M7512 ThermoFisher Scientific) or with anti-TOM20 Rabbit mAb (Cell Signaling Technology #42406) and counterstained using Cy3 conjugated donkey anti-rabbit antibody (Jackson Immunoresearch Laboratories #711-165-152). Nuclei were labelled using Vectashield antifade medium with DAPI (Vector laboratories Inc. #H-1200-10). Confocal pictures were taken at an Airyscan LSM880 (Zeiss) with an 40x Oil objective.

#### Colocalisation experiment in C. elegans

*C. elegans* expressing the MTF and BirA::mCherry under the control of the *myo-3* promoter were incubated wit 5 µM MitoTracker™ Red CMXRos (Thermo Fischer) in M9 (21.6 mM Na_2_HPO_4_, 22 mM KH_2_PO_4_, 8.5 mM NaCl, 18.7 mM NH_4_Cl) supplemented with 2.5% OP50 *E. coli* culture for 22 h at 20°C in the dark. Nematodes were anaesthetised with 250 mM sodium azide and mounted onto a slide with a 3% agarose pad. The slides were covered with a coverslip and image acquisition was the same as described above for *A. thaliana*.

#### For movie

Seven-day old *Arabidopsis* seedlings were placed in a drop of 10 % glycerol on a microscopic coverslip (24 × 50 mm) covered with a coverslip (18 × 18 mm). Time series were collected using the LSM 880 Indimo, AxioObserver (Zeiss) with the Plan-Apochromat 40x/1.4 Oil DIC M27 objective, the MBS 488 beam splitter and excitation wavelength of 488 nm, 1.0% and detection wavelength 411-695 nm. The pixel-dwell was 2.10 µs.

#### Confocal microscopy of bead bound mitochondria

After isolating mitochondria using mRACE for seedlings, 5µL of the samples were placed on a microscope slide and covered with a cover slide. Pictures were taken using the LSM 880 Indimo, AxioObserver (Zeiss) with the Plan-Apochromat 40x/1.4 Oil DIC M27 objective, the MBS 488 beam splitter and with excitation wavelength of 488 nm, 2.0%. The pixel-dwell was 9.31 µs.

#### Confocal microscopy on cp_YFP targeted mitochondria

Microscopy was performed using Zeiss CLSM 880 Confocal Laser Scanning Microscope with excitation wavelength of 488 nm, 2.0 %. Detection wavelength was 406-604 nm to get maximum signal. Imaging was done using Plan-Apochromat 20x/0.80 M27 lens. Time series were collected with time intervals of 1.5 secs. Quantification of the fluorescence intensities was done using ImageJ and graphs were prepared in Excel.

### LC-MS/MS-based quantitative proteome analyses

Proteins were reduced, alkylated and digested on the pulldown beads or in solution for samples originating from gradient isolations. Further sample processing and LC-MS/MS data acquisition were performed as described previously (45). LC-MS/MS analysis was performed by using an EASY-nLC 1200 (Thermo Fisher) coupled to a Q Exactive HF mass spectrometer (Thermo Fisher). Separation of peptides was performed on 17 cm frit-less silica emitters (New Objective, 0.75 µm inner diameter), packed in-house with reversed-phase ReproSil-Pur C_18_ AQ 1.9 µm resin (Dr. Maisch). The column was constantly kept at 50 °C. Peptides were eluted in 115 min applying a segmented linear gradient of 0 % to 98 % solvent B (solvent A 0 % ACN, 0.1 % FA; solvent B 80 % ACN, 0.1 % FA) at a flow-rate of 300 nL/min. Mass spectra were acquired in data-dependent acquisition mode according to a TOP15 method. MS spectra were collected by the Orbitrap analyzer with a mass range of 300 to 1759 m/z at a resolution of 60,000 FWHM, maximum IT of 55 ms and a target value of 3×10^6^ ions. Precursors were selected with an isolation window of 1.3 m/z, and HCD fragmentation was performed at a normalized collision energy of 25. MS/MS spectra were acquired with a target value of 10^5^ ions at a resolution of 15,000 FWHM, maximum injection time of 55 ms and a fixed first mass of m/z 100. Peptides with a charge of +1, > 6, or with unassigned charge state were excluded from fragmentation for MS^2^, dynamic exclusion for 30 s prevented repeated selection of precursors.

Processing of raw data was performed using the MaxQuant software version 1.6.17.0 (46). MS/MS spectra were assigned to the Araport11 protein database. During the search, sequences of 248 common contaminant proteins as well as decoy sequences were automatically added. Trypsin specificity was required and a maximum of two missed cleavages was allowed. Carbamidomethylation of cysteine residues was set as fixed, oxidation of methionine, deamidation and protein N-terminal acetylation as variable modifications. A false discovery rate of 1 % for peptide spectrum matches and proteins was applied. Match between runs, as well as LFQ and iBAQ quantification were enabled.

MaxQuant output tables were further processed in R (47). Potential contaminants and reverse hits were removed. Protein locations were extracted from the SUBAcon4 database (26). Plots were done with the ggplot2 package (48).

## Data availability

MS raw data is available under the following link for review (49) and will be made publicly available upon publication of the article.URL

https://repository.jpostdb.org/preview/113242714060d2fd355f451

Access key

8994

## Acknowledgments

We sincerely thank Thomas Schwarz for his insightful and valuable inputs in this manuscript. We thank Holger Doschke and Dario Furlani for sharing their knowledge and expertise on AFM and confocal imaging respectively. We also thank Markus Schwarzländer for kindly providing the mitochondrial-targeted cpYFP expressing *Arabidopsis thaliana*, Col-0 seeds (*p35s::mt-cpYFP+/+*).

We gratefully acknowledge financial support from the European Research Council to RG (ERC Consolidator Grant “bi-BLOCK” ID. 646644, ERC Proof of Concept Grant “TriVolve” ID 957547), funding by the Deutsche Forschungsgemeinschaft to IF (INST 211/744-1 FUGG) to RG (637149), to KM (MA4172/12-1 & MA 4172/15-1) and to JK (DFG KI1988/5-1).

