## Supplementary data for "A versatile mitochondria isolation- and analysis-pipeline generates 3D nano-topographies and mechano-physical surface maps of single organelles"

### Supplementary figures:

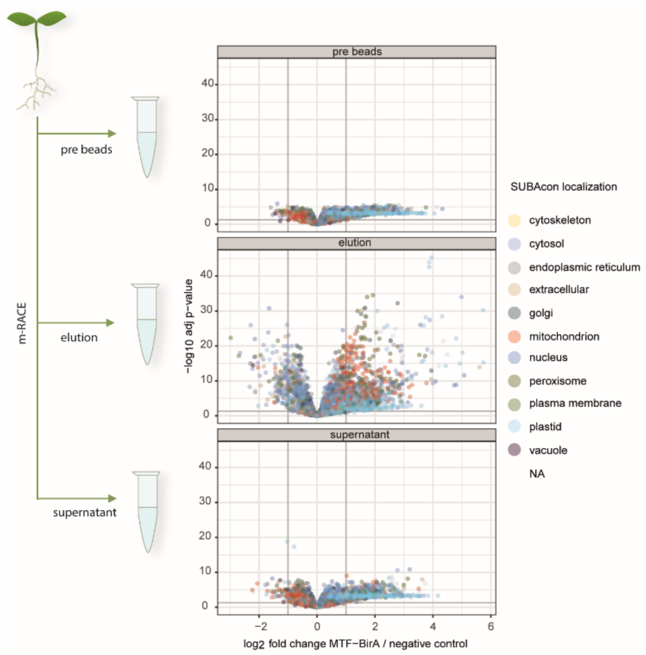


**Supplementary Figure 1: Enrichment of mitochondrial proteins during mRACE.** Samples of 100 µL were collected at the pre beads step and from the supernatant in addition to the 100 µL eluate and compared to samples from plants not expressing the mRACE constructs (negative control). While neither pre beads nor supernatant samples show an enrichment of mitochondrial proteins there is a significant increase of mitochondrial proteins from all compartments in the eluate.

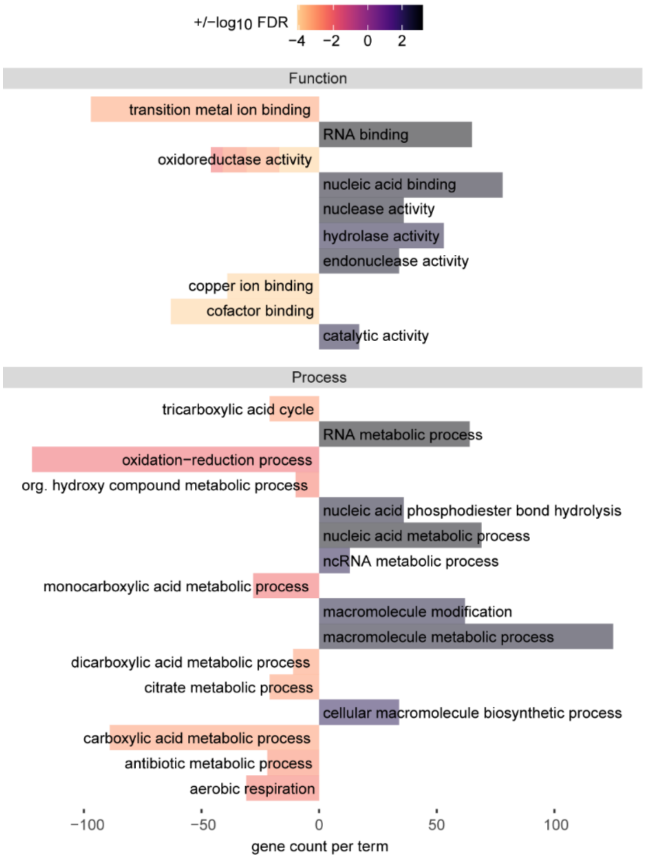


**Supplementary Figure 2:** Box plots correlating the number of tissue specifically enriched proteins per GO term in either tissue. Number of proteins per term depicted on the x-axis. Negative values indicating an enrichment in roots (left), positive in seedlings (right). The respective log10 FDR is used as fill colour of the respective bar. The sign is indicating the enrichment in root (-) or seedling (+) samples.

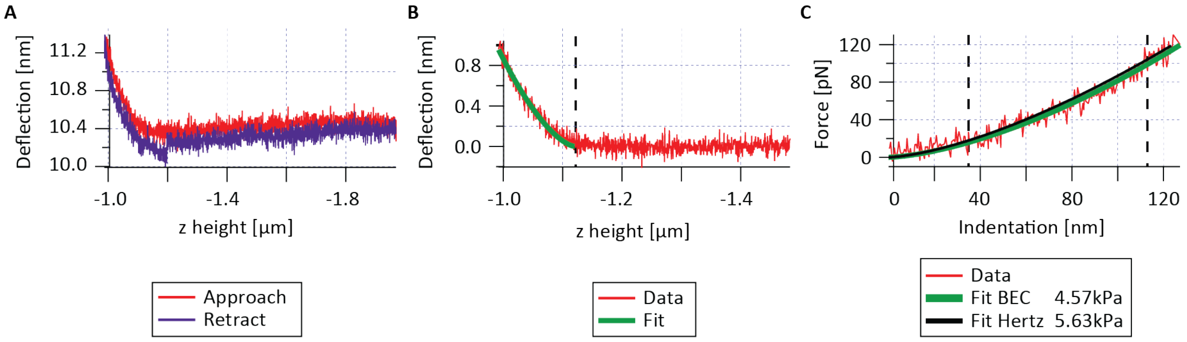


**Supplementary Figure 3: Raw data processing on a typical force curve with Hertz and BEC fit:** Panel A shows a typical force curve on a mitochondrion showing the deflection data vs z height as recorded during approach and retract. The approach data was analysed after subtracting tilt of the force curve in the non-contact part and removing data points off the surface (Panel B), which can be seen from the smaller z-range in B compared to A. From these data we calculated the indentation and force (Panel C) and fitted those data with the appropriate model from contact mechanics. Conventionally the Hertz model (here for the case of a spherical indenter with 75 nm radius) is used (black fit line in C). Even though the indentation (maximum ~ 130 nm) is small compared to thickness of the sample (at this force curve: 732 nm) using the bottom effect correction (BEC) from Garcia et al, we obtained a substantially different Young's modulus: 4.57 kPa for BEC instead of 5.63 kPa for a conventional Hertz fit. The fitted force vs indentation curves are denoted in Panel C in green for BEC and black for Hertz. The black dashed vertical lines in C denote the range if data which has been used for the fit. As can be seen from the dashed vertical line in B the contact point is at a z height of -1.116 µm.

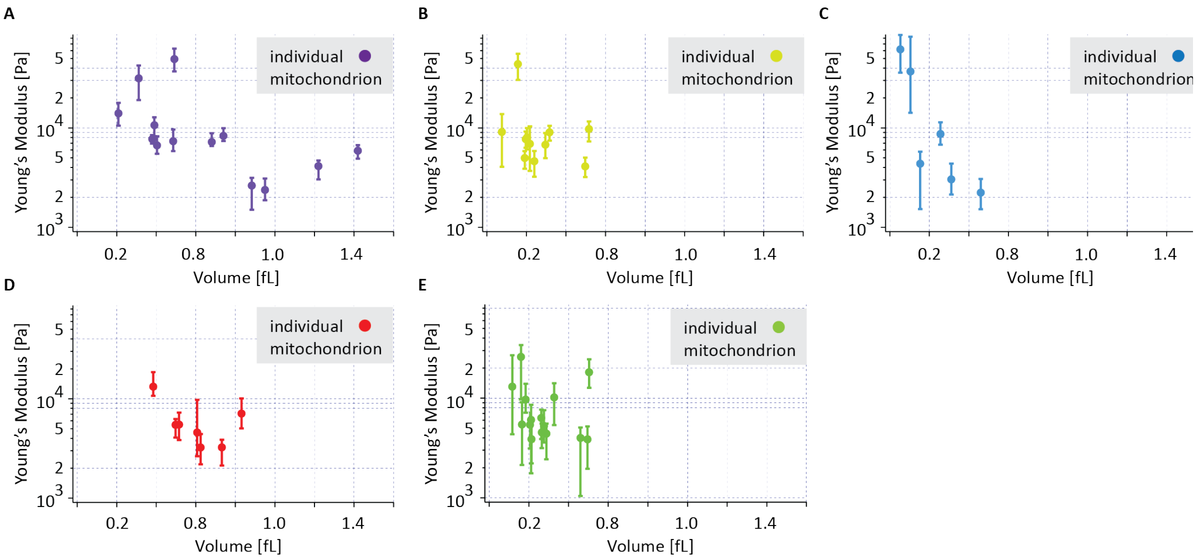


**Supplementary Figure 4:** Figure 4F is compilation of 5 independent experiments which are individually depicted here. Whiskers represent 25th and 75th of width distribution of YM values.
